# Life-history and connectivity shape the viromes of a migratory and river-resident galaxiid

**DOI:** 10.64898/2026.10.08.757744

**Authors:** Ryan R. Easton, Stephanie Waller, Hayley Bosley, Isa de Vries, Matt Jarvis, Jeremy Dubrulle, James Hodgkinson-Bean, Gerard P. Closs, Jemma L. Geoghegan

## Abstract

Migration can increase population connectivity and gene flow but may also facilitate pathogen transmission among hosts and habitats. The way life-history strategies shape viral diversity in closely related freshwater fishes remains poorly understood. Here, we used metatranscriptomics to compare the viromes of two fish in the family Galaxiidae with contrasting life-history strategies: amphidromous īnanga (*Galaxias maculatus*) and river-resident Gollum galaxias (*Galaxias gollumoides*). We analysed 54 pooled libraries derived from 416 individuals collected across 27 sites spanning multiple catchments and fish-passage barriers in southern New Zealand. A total of 13 novel fish-associated viruses were identified from five viral families (*Astroviridae*, *Circoviridae*, *Coronaviridae*, *Hepeviridae,* and *Picornaviridae*). Virome composition differed markedly between species, with only a single virus-positive pool detected among īnanga, whereas Gollum galaxias harboured 11 novel viruses, with circoviruses dominating the virome. No fish-associated viruses were detected in post-larval juveniles of either species, and alpha diversity did not differ significantly among catchments, sites or positions relative to fish-passage barriers. However, beta diversity analyses revealed significant spatial structuring of viral communities among sites and catchments, indicating localised variation in virome composition. Contrary to our predictions, the migratory species exhibited substantially lower viral diversity than the river-resident species. The absence of detectable viruses in juveniles, combined with the greater viral richness of the longer-lived Gollum galaxias, suggests that host longevity and cumulative environmental exposure may be important drivers of virome assembly. These findings demonstrate that host life-history strategy influences virome composition and reveal unexpectedly low viral diversity in a globally widespread migratory galaxiid, providing new insights into virus-host ecology and connectivity in freshwater fishes.

## Introduction

Migration has numerous benefits for organisms, including enabling access to higher quality or more abundant food, habitat, and other resources outside their natal area (Alerstam & Bäckman, 2018; Brönmark et al., 2008; Fudickar et al., 2021; Lennox et al., 2016). Furthermore, migrations facilitate the movement of individuals, larvae, and genes between populations, increasing the connectivity of otherwise unconnected groups (Dingle & Drake, 2007; Lowe & Allendorf, 2010; Marra et al., 2006; Webster et al., 2002). While increased connectivity can provide demographic and genetic benefits at the population level, migration may impose costs on individuals through increased exposure to parasites and pathogens encountered along migratory pathways and at aggregation sites, potentially reducing survival and fitness (Breyta et al., 2022a; Krkosek et al., 2007; Miller et al., 2014; Poulin & de Angeli Dutra, 2021; Vollset et al., 2021).

Varying life-history strategies have been shown to shape viral transmission dynamics and disease expression (Altizer et al., 2011; Miller et al., 2014; Peacock et al., 2020). Migratory pathways therefore represent potential routes for disease and viral transmission between hosts and transfer between habitats (Krkosek et al., 2009; Poulin et al., 2012; Vollset et al., 2021). Disease transmission of parasites, viruses, and bacteria has been described in migratory salmonids, particularly in relation to migratory pathways that encounter aquaculture operations, and the transition periods between freshwater and marine habitats (Krkosek et al., 2009; Vollset et al., 2021). What is lacking however is the investigation into how the viromes of closely related (congeneric) migratory and river-resident populations differ.

Populations of river-resident fish species, and particularly those in headwater streams, are among the most ecologically and genetically isolated fish populations globally (Fagan, 2002; Gido et al., 2015; Zarri et al., 2022), yet many trace back to migratory species that lost the need or ability to migrate (Delgado & Ruzzante, 2020; Dodson et al., 2013; Kendall et al., 2015; Waters & Wallis, 2001). The degree of population connectivity and genetic isolation likely shapes their virome, with the expectation that there will be similarities in viromes between migratory and river-resident populations, but that the more isolated populations will exhibit the loss of viruses retained in the migratory mixing populations.

An exceptional group of fish to use to test this theory is the family Galaxiidae, found in the Southern Hemisphere, which has populations of diadromous and river-resident species, some of which overlap in distribution across their respective ranges. Aotearoa New Zealand has five species of diadromous (amphidromous) galaxiids (three endemic) (Department of Conservation, n.d.-a), and 12 formally recognized, with a further 13 indeterminate taxa of river-resident galaxiids (Department of Conservation, n.d.-b). These river-resident non-migratory species are all likely derived from a single common migratory ancestor that lost diadromy, and then speciated via isolation through geological processes (Campbell et al., 2022; Carrea et al., 2014; Crow et al., 2009; Waters et al., 2015). While river-resident species are genetically isolated from other conspecifics/congenerics, there is increasing evidence that diadromous species are also isolated from each other at the population scale (Astorga et al., 2022; Augspurger et al., 2023).

The virosphere of freshwater fish on the three main islands of New Zealand is relatively understudied, with the majority of the migratory-galaxiid virome knowledge coming from the Chatham islands (Grimwood et al., 2023). Other widespread species with sympatric distributions have been investigated, including two freshwater eels (*Anguilla spp.*) (Waller et al., 2024), *Retropinna retropinna* (common smelt), *Gobiomorphus cotidianus* (common bully) (Perry et al., 2022), and southern pouched lamprey (*Geotria australis*) (Darnley et al., 2026; Miller et al., 2021).

Here, we examine the viromes of two congeneric fish species in the family Galaxiidae (genus *Galaxias*) resident to the Murihiku (Southland) region of Te Waipounamu (South Island) Aotearoa, New Zealand, using a total RNA metatranscriptomic approach. First, īnanga (*Galaxias maculatus*), one of five amphidromous migratory galaxiids in New Zealand and one of the most widely distributed freshwater fishes in the world (Waters et al., 2000). They are predominantly annual species (McDowall et al., 1994; Watson et al., 2021) and are the most common galaxiid in New Zealand, comprising most of the economically important commercial and recreational whitebait fisheries (Yungnickel et al., 2020). Second, Gollum galaxias (*Galaxias gollumoides*), a longer-lived (4-7 years; (Fenton, 2022)) non-migratory river-resident species of ‘roundhead’ galaxiid common throughout the Southland region. Populations of Gollum galaxiids are geographically isolated from conspecifics within smaller tributaries of larger river systems, resulting in limited downstream connectivity and increasing genetic isolation (Crow et al., 2009; Waters et al., 2015). While the Gollum galaxiid is not directly descended from īnanga (Burridge et al., 2012; Waters et al., 2010), the distribution of these two species does occasionally overlap in lower altitude coastal catchments although they are mostly allopatric (Leathwick et al., 2008).

Our aim is to investigate whether the viromes of these two species could be used to assess population connectivity across the Southland region, and if the loss of river connectivity through fish passage barriers has had an impact on virome connectivity. To assess these patterns, we explored the viral richness and abundance of these two species across sites within and between multiple catchments, different life-history stages, and capture location upstream/downstream of fish passage barriers. We hypothesise that īnanga will show greater virome similarity across the region than Gollum galaxiids, given their amphidromous life cycle, which enables greater population mixing than that of Gollum galaxiids. In contrast, most populations of Gollum galaxiid are geographically and functionally isolated with a distribution that is largely restricted to small headwater streams and upstream of most populations of īnanga (Waters et al., 2015).

## Methods

### Sample locations

Sites were located within the Murihiku (Southland) region of Te Waipounamu (South Island), New Zealand (Figure 1a). Sites were selected based on the presence of a potential partial or complete barrier to fish passage and distributed across the region to capture variability across the distribution of each species within and between catchments and sites. Adult īnanga and Gollum galaxiids, as well as post-larval juveniles of each species, when possible, were opportunistically collected within a 150-meter reach upstream and downstream of each barrier location. This sampling design created a spatial hierarchical structure with individuals of each species nested by site, life history stage, upstream/downstream collection location of each fish passage barrier, and catchment.

**Figure 1.**
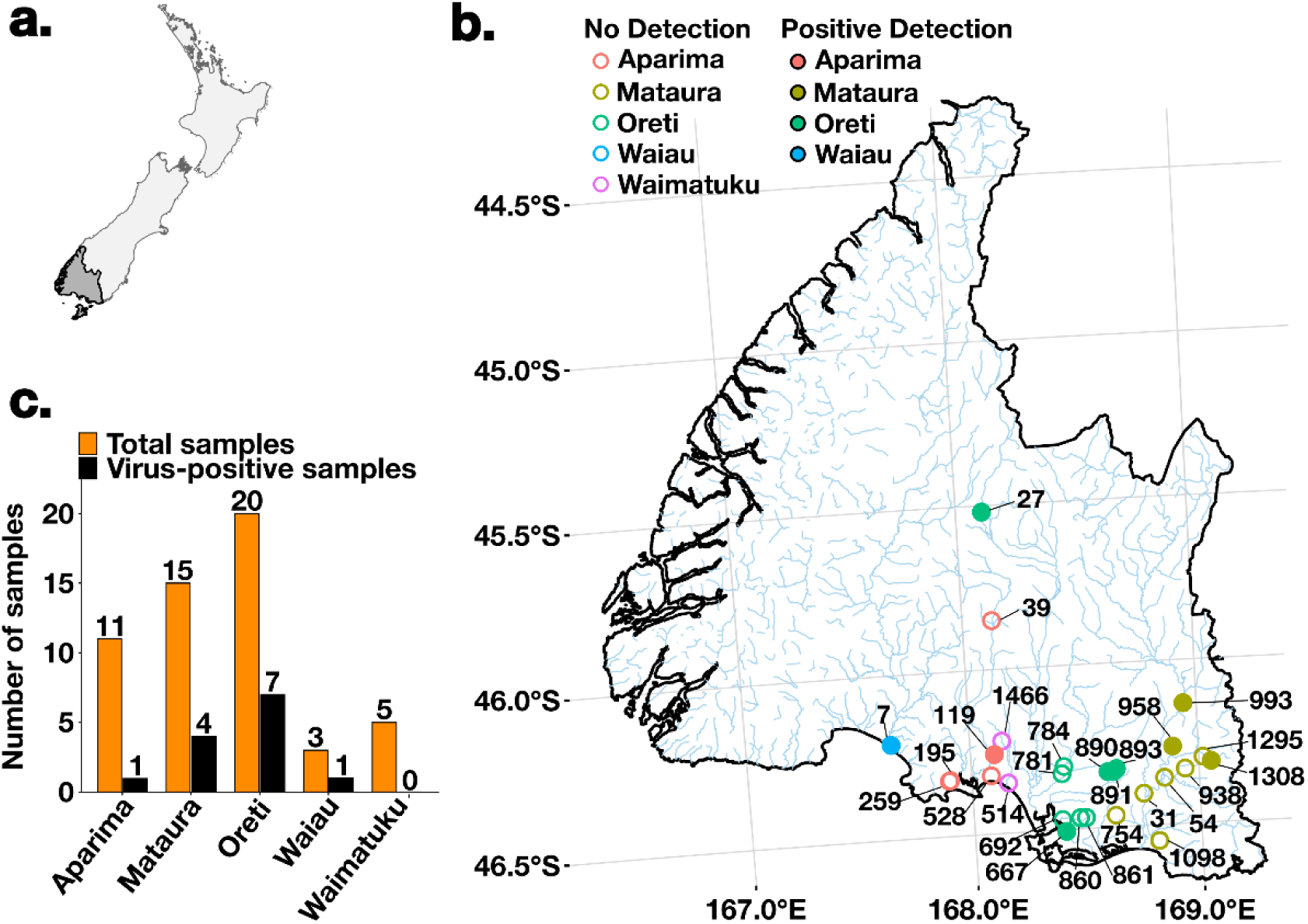
Map of sampling locations with viral detection by catchment. (a) Map of New Zealand with the Murihiku (Southland) Region of Te Waipounamu (South Island) highlighted in grey. (b) Map of the 27 sampling locations (identified by location number), coloured by catchment area, across the Southland Region. Closed circles denote positive viral detection, and open circles denote zero viral detection. (c) Total number of sites sampled by catchment (orange) with the total number of virus-positive samples detected in each catchment (black).

### Liver and gill tissue collection

Between September 2022 and April 2023, 235 īnanga and 205 Gollum galaxiids were collected across 27 sites in the Southland Region of New Zealand’s South Island (Figure 1b). Fish were collected by electrofishing, conducted using a Smith-Root® LR-24 backpack machine (Smith Root Inc., USA), set between 200 – 300 V with a frequency of 30 Hz (equivalent to a 4ms pulse width) with the aim of collecting 10 fish from each site where possible. Captured fish were euthanised with an overdose of AQUI-S (University of Otago Animal Use Protocol 21-156), and their gills and livers were immediately dissected with a sterile scalpel. Tissue samples were submerged in 1 mL of RNA stabilisation solution (DNA/RNA Shield, Zymo Research) and stored at 4°C until they were transferred to the University of Otago, Dunedin, New Zealand where they were stored at −80°C until total RNA was extracted. More information regarding the sample collection date, location, and number of individuals sampled per site is provided in Supplementary Table 1.

### Extraction of total RNA from fish livers and gills

Frozen tissue samples stored in DNA/RNA Shield were thawed, and the tissue was placed in 15-mL RNase-free round-bottom tubes containing lysis buffer. The samples were homogenised for one minute using a TissueRuptor (QIAGEN). Total RNA was extracted using the RNeasy Plus Mini Kit (QIAGEN) according to the manufacturer’s protocol, with minor alterations. Briefly, two ethanol wash steps were added to remove any residual guanidine contamination. Extracted RNA was quantified using a NanoDrop spectrophotometer. RNA was obtained from 212 of the 235 īnanga and 204 of the 205 Gollum galaxiids at concentrations suitable for downstream processing. Equal volumes of RNA eluate from 1-10 individuals were pooled into 54 libraries based on the sampling location and life stage, post-larval juvenile, or adult (Supplementary Table 1).

### RNA sequencing

Extracted RNA was subject to total RNA sequencing. Libraries were prepared using a Stranded Total RNA Prep with Ribo-Zero Gold Kit (Illumina). Paired-end 150-bp sequencing of the RNA libraries was performed on the Illumina NovaSeq X platform.

### Virome assembly and virus identification

Sequencing reads were quality-trimmed using Trimmomatic v0.38 (Bolger et al., 2014). Adaptor sequences were removed, low-quality bases with quality scores of less than 3 were trimmed from the 5′ and 3′ ends, and sliding window trimming (4 bases, average quality <5) was applied. Reads shorter than 25 nucleotides after trimming were discarded and only paired reads retained after trimming were used for downstream analyses. Following quality control, reads were de novo assembled into contigs using MEGAHIT v1.2.9 (Li et al., 2015). Contigs were then compared against the NCBI nucleotide (nt) and non-redundant protein (nr) databases using BLASTn v2.15.0 (Altschul et al., 1990) and DIAMOND v2.1.9 (Buchfink et al., 2015) to identify viral sequences. A similarity threshold of 1×10 for the nt database and 1×10 ¹ for the nr database was applied to reduce false positive results.

### Estimating the abundance of viral sequences

Virus abundance was estimated by mapping reads back to the assembled contigs using Bowtie2 v2.4.4 (Langmead & Salzberg, 2012) and SAMtools v1.9 (H. Li et al., 2009). Sequences representing <0.1% of the read counts in another library and sharing >99% nucleotide identity were considered likely index-hopping artifacts and excluded from further analysis (Chang et al., 2024). Total viral abundance estimates for viruses across viral families and orders were compiled across all libraries. Estimated abundances per million reads were standardised to the number of paired reads per library.

### Phylogenetic analysis

Translated conserved viral protein sequences (i.e., RNA dependent RNA polymerase (RdRp), DNA polymerase) were aligned with representative protein sequences from the same virus family obtained from NCBI RefSeq as well as the closest BLASTp hits using MAFFT v7.490 (Katoh & Standley, 2013). Ambiguously aligned regions were removed using trimAL v1.2rev59 (Capella-Gutiérrez et al., 2009) with the gap threshold flag set to 0.9. Phylogenetic trees for each viral species/family/order were then estimated using the maximum likelihood method in IQ-TREE web server (Trifinopoulos et al., 2016), employing the LG amino acid substitution model with 1000 ultra-fast bootstrapping replicates. The resulting phylogenetic trees were annotated using Figtree v1.4.4 (Rambaut, 2018). Phylogenetic comparison with known viruses was used to infer the probable host origin of each virus. Specifically, viruses that were phylogenetically distinct from vertebrate host viruses were assumed to be more likely associated with diet, microbiome or environmental sources. Viruses that clustered with known fish viruses were subject to further evolutionary analysis.

### Viral abundance, diversity and network analysis

All plots were created in R v4.5.2 using RStudio v2024.04 with the tidyverse ggplot2 package (Villanueva & Chen, 2019). For each library, raw read counts were first standardised as reads per million (RPM), calculated as the number of reads mapping to a viral family divided by the total number of reads in that library, multiplied by one million. RPM values were further normalised by the total RPM within each library so that the values summed to one, allowing direct comparison across samples of differing sequencing depths.

To examine what shaped viral diversity and connectivity across the region, we compared viral richness and alpha diversity across species, sites, catchments, and upstream/downstream of fish passage barriers. As no viruses were identified in post-larval juveniles, life-history stage was not analysed. Total length and visual developmental stage identification was used to distinguish juveniles from adults (Fenton, 2022). Īnanga are visually distinguishable between the juvenile “whitebait” stage and adults. Gollum galaxiids mature at age 0+ (Dunn et al., 2022), typically growing up to 70 mm in length but can reach 140 mm (Fenton, 2022). Detailed length at age information is unavailable, therefore we investigated the length distribution of “adult” and “juvenile” to ensure that the length classification we used to distinguish them was in-line with the limited literature available, with both individual lengths (p = 2.56 × 10 ¹²) and pooled lengths (p = 0.0062) were significantly different from each other. Viral richness counts the number of taxa present, and alpha diversity used the Shannon diversity index, which measures richness and evenness calculated from viral family-level standardised abundance data (RPM) using the Vegan package (Oksanen et al., 2001). Differences across paired comparisons of independent groups, species, and barrier location (upstream/downstream) were evaluated using a non-parametric Wilcoxon rank-sum test, whereas sites and catchments were evaluated using a Kruskal–Wallis test. When significant differences were detected, Dunn’s post hoc test with Benjamini–Hochberg correction for multiple comparisons was applied. Boxplots of richness and Shannon diversity were generated using ggplot2.

Beta diversity was assessed using Bray-Curtis dissimilarity, calculated from family-level viral abundance values (RPM), to evaluate how species, site, catchment, and barrier location (upstream/downstream) influenced the overall viral community composition. A Principal Coordinates Analysis (PCoA) was performed to visualise the differences in viral community composition between sample sites, as it accepts non-Euclidean distance measures like Bray-Curtis. Ordinations were created using ggplot2 and coloured by site, catchment, species, and barrier location. To statistically test the differences in viral composition between species, sites, catchments, and barrier locations (upstream/downstream), an analysis of similarities (ANOSIM) was performed using the Bray-Curtis distances with 9999 permutations. ANOSIM was used because it is robust to zero-inflated, low-richness data with unequal group sizes and sparse detections.

To visualise the viral community composition by species and life-history stage, we constructed a heatmap of viral family abundance across libraries. We tested theRPM values were normalised by the total RPM within each library so that the values summed to one, allowing a direct comparison across samples of differing sequencing depths. Relative viral family abundance was plotted using a heatmap in the ggplot2 package (Villanueva & Chen, 2019). Relative abundances are represented on a log10 scale. Libraries were ordered by site and species to highlight ecological structure.

To investigate the relationships between viral families shared between and among species, a bipartite network analysis was created using the ggbipart package (Jordano, 2026) within the ggplot2 (Villanueva & Chen, 2019) environment. To quantify the structural properties of the network, nestedness was calculated using the bipartite package in R (Dormann et al., 2020). Nestedness describes the degree to which species with fewer viral associations harbour subsets of those found in more virus-rich hosts.

## Results

Total RNA was extracted from combined gill and liver tissues from 212 īnanga combined into 28 pools (15 sample locations) and 204 Gollum galaxiids combined into 26 pools (16 sample locations) from the Murihiku Southland region (Figure 1, Supplementary Table 1). Pools were based on site location, life-history stage, and position upstream/downstream of a fish passage barrier, with each species overlapping at four sites (five Gollum galaxiid pools and six īnanga pools). Metatranscriptomic sequencing generated between ∼63.7 and 105.3 million paired-end reads per library (Supplementary Table 1).

Analysis of the metatranscriptomes revealed the sequences of two novel īnanga viruses and 11 novel Gollum galaxiid viruses from five different viral families across the 54 pools (Figure 2a-c, Supplementary Table 2). Of the 28 īnanga pools, only one had fish-associated viral sequences from the family *Coronaviridae*. The family *Circoviridae* was the most frequently occurring viral family, identified in eight of the 26 Gollum galaxiid pools (Figure 2c). Only one pool of Gollum galaxiids had more than one fish-associated virus present, located in the Mataura catchment (Figure 2c).

**Figure 2.**
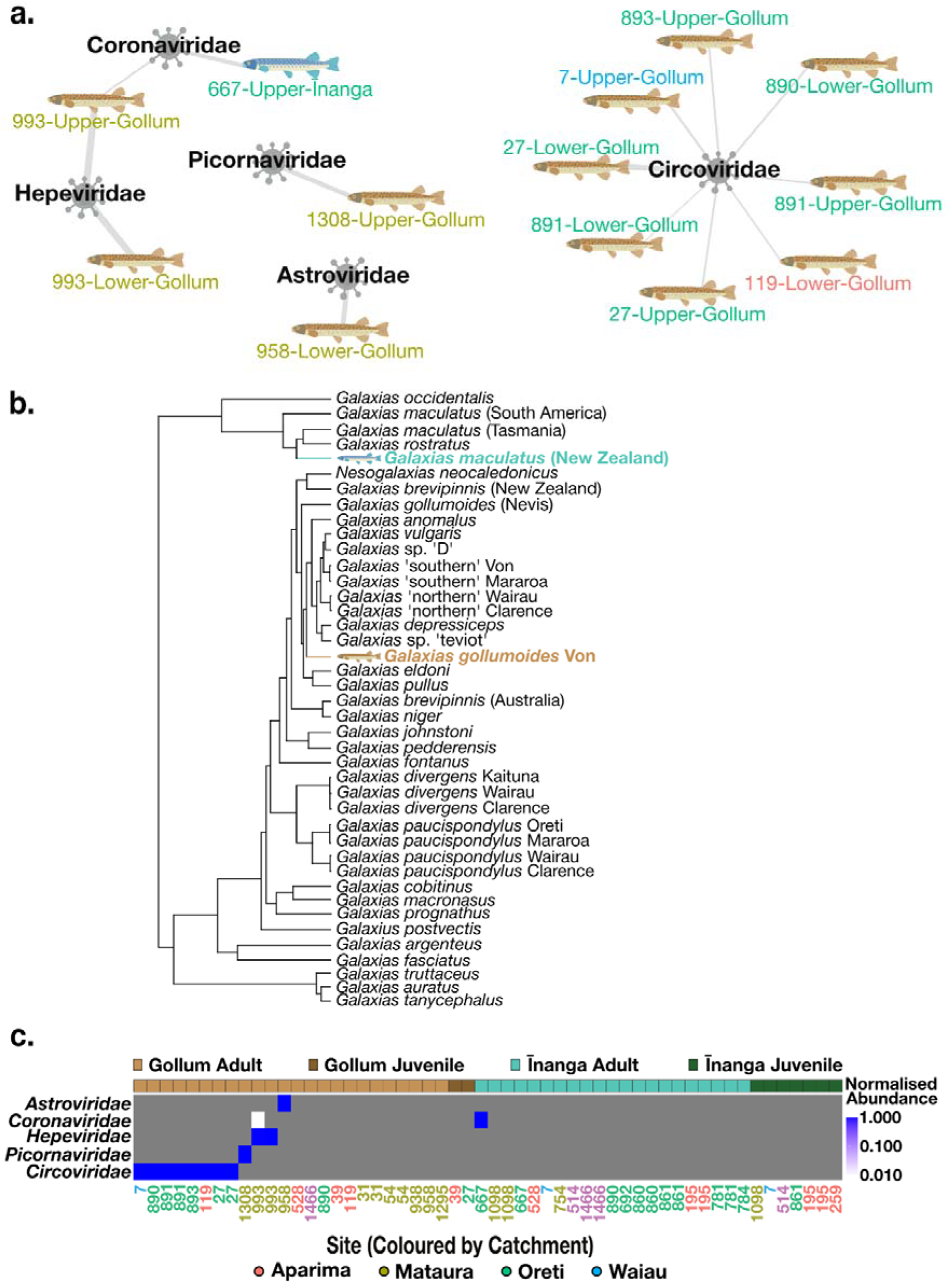
Host-virus associations visualised through a bipartite network and normalized relative abundance heatmap. (a) Bipartite network displaying the association of each viral family between īnanga and Gollum galaxiids. Each node represents a viral family, with line weights derived from the summed reads per million (RPM) values across libraries for each species coloured by catchment. (b) Phylogenetic tree of the relationship between īnanga and Gollum galaxiids (modified from Burridge et al., 2012). (c) Heatmap showing the normalized relative abundance of each viral family by species and life-history stage. Colour intensity indicates relative abundance on a log10 scale, ranging from low (white) to high (blue). Site capture locations below heatmap are coloured by catchment.

### (+)ssRNA viruses

#### Astroviridae

A novel astrovirus was identified in a pool of Gollum galaxiids from the Mataura catchment (Figure 3a and Supplementary Table 2). The partial sequence containing the RdRp shared 47.55% amino acid (aa) sequence identity with European seabass (*Dicentrarchus labrax*) virus (UZX50331.1), previously identified in aquacultured European seabass which were experiencing high fingerling mortality (Louboutin et al., 2026).

**Figure 3.**
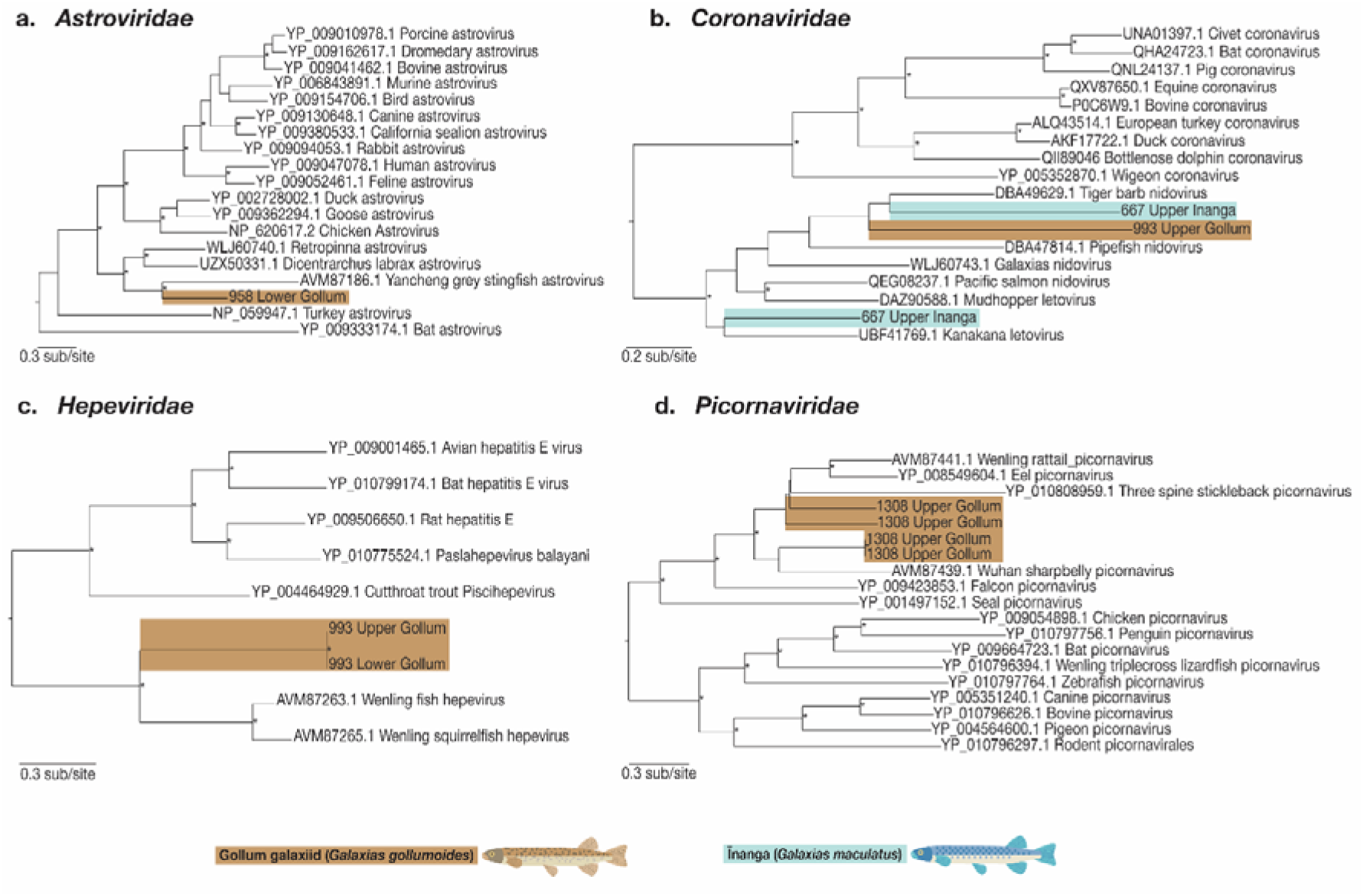
Phylogenetic tree of (+ssRNA) viruses. Maximum likelihood phylogenetic trees were constructed from representative viral transcripts containing RdRp sequences from the families (a) *Astroviridae,* (b) *Coronaviridae,* (c) *Hepeviridae, and* (d) *Picornaviridae*. Viruses identified in this study can be identified by their tip labels indicating site location and species (Gollum galaxiids in brown and īnanga in blue). Branch lengths are proportional to the number of amino acid substitutions per site, with all trees being midpoint-rooted. Main nodes with ultrafast bootstrap values >70% are indicated by an asterisk.

#### Coronaviridae

Two sequences within the family *Coronaviridae* were identified in īnanga and shared <33% amino acid sequence identity and were phylogenetically divergent; therefore, they were determined to represent two viral species (Figure 3b, Supplementary Table 2). The closest BLAST hits were the Tiger barb (*Puntigrus tetrazona*) nidovirus (DBA49629.1) and Broadnosed pipefish (*Syngnathus typhle*) nidovirus (DBA47814) (Buck et al., 2024). A single sequence identified from a pool of Gollum galaxiids identified in a different catchment to the īnanga had a top BLAST hit to an Orange clownfish (*Amphiprion percula*) (DBA48395.1) nidovirus (Buck et al., 2024).

#### Hepeviridae

Two sequences within the family *Hepeviridae* were identified at the same site in Gollum galaxiids, one upstream and one downstream of the fish passage barrier. They are likely the same species, as they had 100% protein alignment, where they overlapped and showed no phylogenetic divergence (Figure 3c, Supplementary Table 2). Each sequence had a 41.23% amino acid sequence identity to the top BLAST hit of Wenling fish hepevirus (AVM87263.1), which was identified from a survey of ray-finned fish (Shi et al., 2018).

#### Picornaviridae

Multiple sequences within the family *Picornaviridae* were identified from a pool of Gollum galaxiids from the Mataura catchment, representing three novel species (Figure 3d; Supplementary Table 2). The partial sequences all had top BLAST hits to fish picornaviruses: 43.45% aa sequence identity with the Wuhan sharpbelly (*Hemiculter leucisculus*) picornavirus (AVM87439.1) (Shi et al., 2018), 51.35% aa sequence identity with a Potamipivirus identified in diseased European eels (*Anguilla Anguilla*) picornavirus 1 (YP_008549605.1) (Fichtner et al., 2013), 55.80% similarity with *Asterropteryx semipunctatus* (Starry goby) picornavirus (UVF58821.1) (Costa et al., 2023), and 44.05% similarity with Wenling rattails (*Coelorinchus multispinulosus*) picornavirus (AVM87441.1) (Shi et al., 2018).

### ssDNA viruses

#### Circoviridae

Multiple partial sequences containing the replication-associated protein region from the family *Circoviridae* were identified in Gollum galaxiids across six sites in three catchments, representing five novel species (Figure 4; Supplementary Table 2). These sequences had top BLAST hits to two circoviruses, and phylogenetically clustered with other distinct fish circoviruses. One set shared 59.32%-60.32% amino acid similarity with and phylogenetically mapped to a novel circovirus (UVF58770.1) identified in Bengal sergeant (*Abudefduf bengalensis*) (Costa et al., 2023). The other group shared 55.71%-68.89% amino acid similarity with a circovirus identified in the feces of Weddell seals (*Leptonychotes weddellii*), which also phylogenetically clustered with other fish circoviruses.

**Figure 4.**
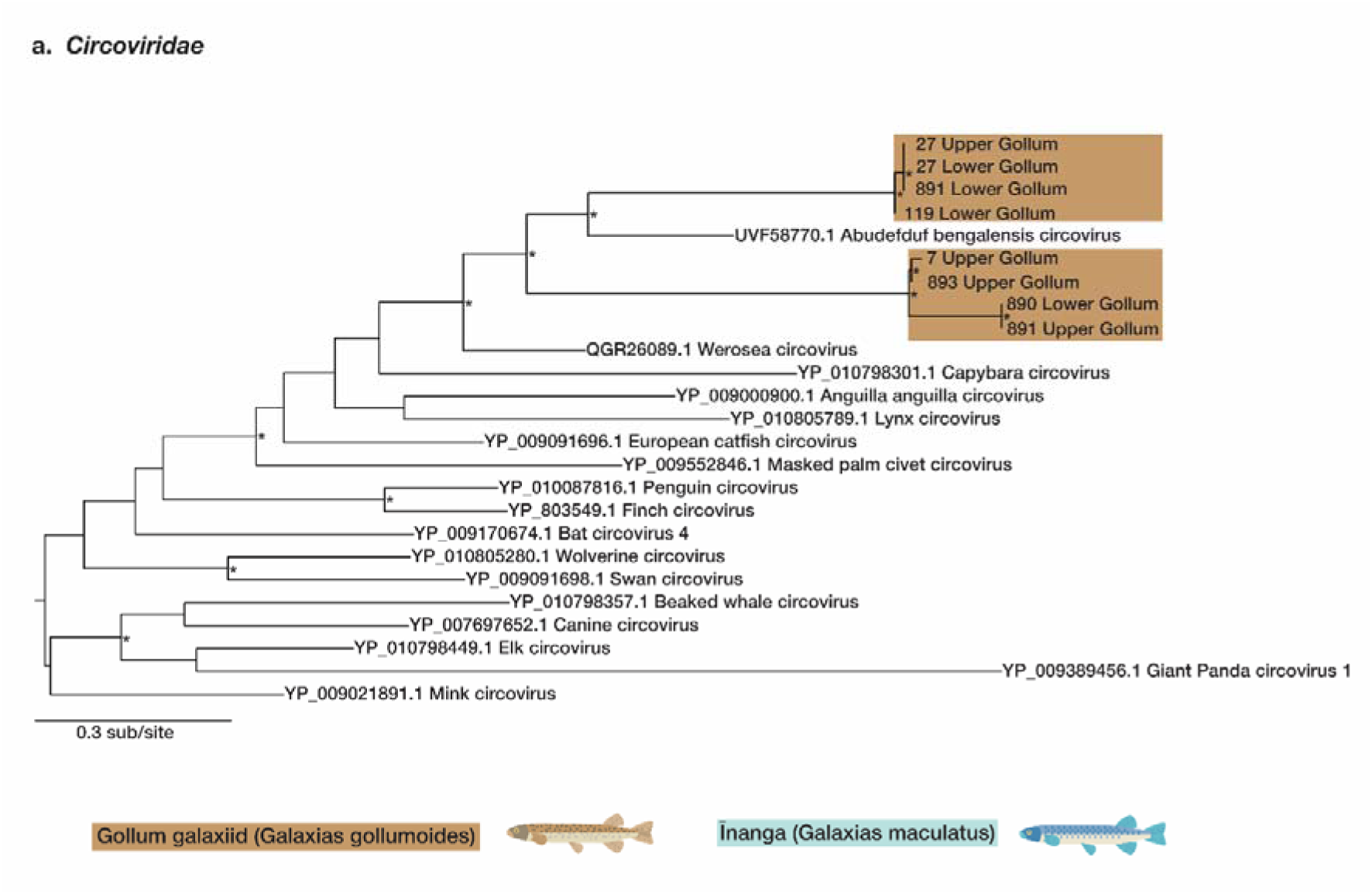
Phylogenetic tree of the DNA virus *Circoviridae*. Maximum likelihood phylogenetic trees were constructed from representative viral transcripts containing the replication-associated protein. Viruses identified in this study can be identified by their tip labels indicating site location and species (Gollum galaxiids in brown and īnanga in blue). Branch lengths are proportional to the number of amino acid substitutions per site, with the tree being midpoint-rooted. Main nodes with ultrafast bootstrap values >70% are indicated by an asterisk.

### Viral abundance and diversity

To investigate the connectivity of viral richness, diversity and distribution in relation to species and landscape (barrier location (upstream/downstream), catchment, and site), we examined virus richness, which measures the number of families with abundance greater than zero. For alpha diversity, the Shannon diversity index was investigated but not reported because the viral abundance was sparse, with most libraries containing no detected fish viruses or only a single family, with only one pool containing two families. Consequently, values were clustered around zero and provided no additional information for our analysis, so are not presented here. Gollum galaxiids had the greatest virome richness of the two species; however, as there was only a single virus-positive īnanga pool, no statistical comparison was performed due to insufficient sample size (Figure 5a). Viral richness did not differ significantly by site (Kruskal–Wallis χ² = 34.84, df = 26, p = 0.12), catchment (Kruskal– Wallis χ² = 4.25, df = 4, p = 0.3727), or between groups upstream and downstream of the identified fish passage barriers (Wilcoxon rank-sum test, p = 0.498). Because īnanga only had a single virus family detected in one pool, we further calculated richness for Gollum galaxiids only. No significant differences were revealed by site (Kruskal–Wallis χ² = 17.08, df = 15, p = 0.3143), catchment (Kruskal–Wallis χ² = 5.327, df = 4, p = 0.2554), or upstream and downstream of the identified fish passage barriers (Wilcoxon rank-sum test, p = 0.3836).

**Figure 5.**
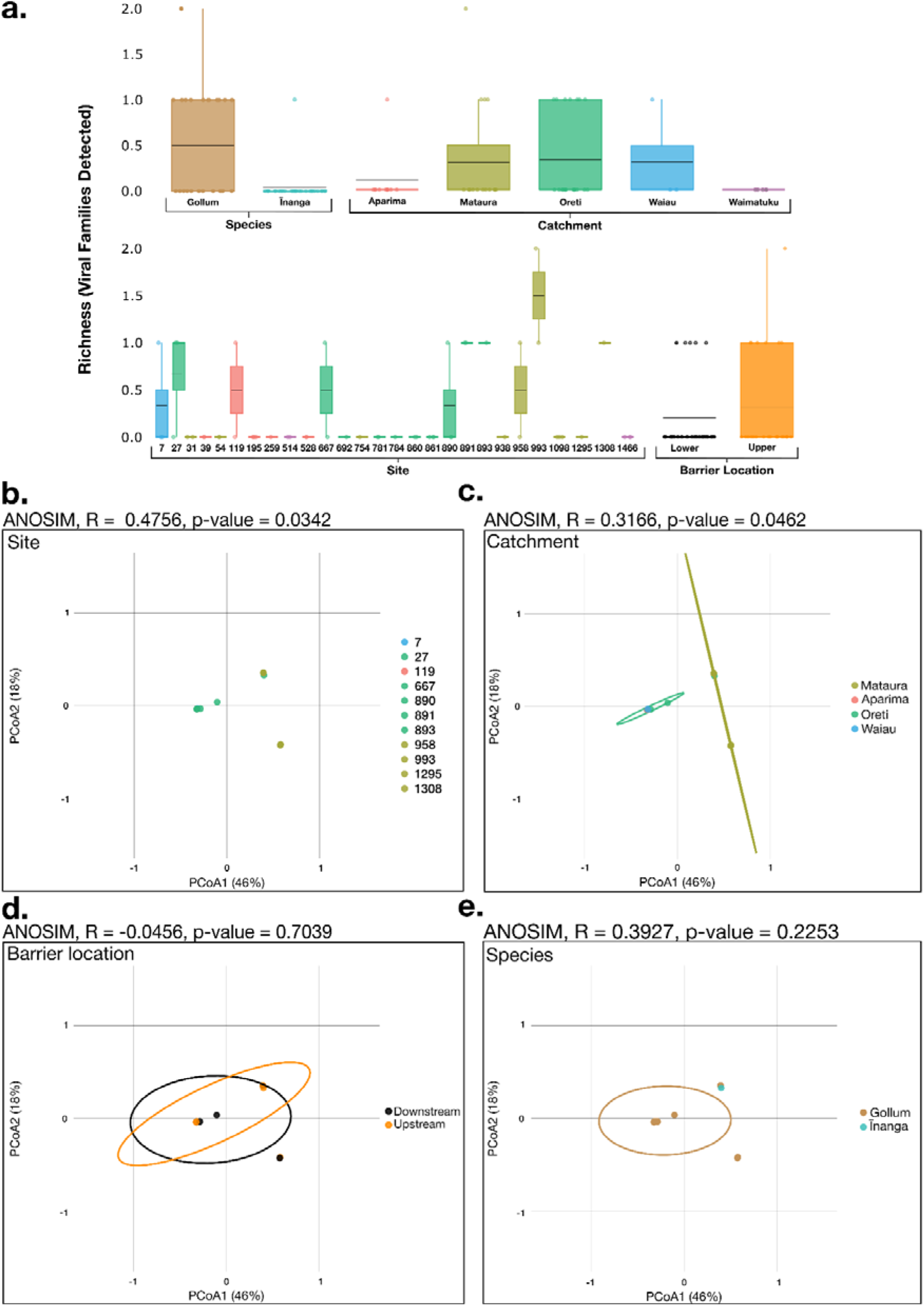
Alpha and beta diversity of īnanga and Gollum galaxiid viromes. (a) Boxplot of viral family richness (alpha diversity) across species, catchment, site, and barrier location (upstream/downstream). PCoA plots of beta diversity across (b) site, (c) catchment, (d) barrier location (upstream/downstream), and (e) species, with associated significance values (R Statistic and p-value) from Analysis of Similarity (ANOSIM) tests above each plot.

To assess how virome composition was influenced across the landscape by site, catchment, and fish passage barriers, we examined the beta diversity of each species, as well as Gollum galaxiids. Beta diversity was visualised using Principal Coordinates Analysis (PCoA), based on Bray–Curtis dissimilarities, with similarity between groups analysed with an Analysis of similarities (ANOSIM) non-parametric test for significance. The first two PCoA axes explained a moderate proportion of the total variation in the viral community composition (PCoA1 46%, PCoA2 18%). At the site level, PCoA suggested a stronger separation among sites, with several sites forming distinct clusters in the ordination space. ANOSIM identified significant differences in viral community composition among sites (ANOSIM R = 0.4756, p = 0.0342, Figure 5b), indicating site-level variations in virome composition. The analysis revealed moderate spatial structuring of the viral community composition among the catchments (Figure 5c). Principal coordinates analysis (PCoA) showed partial clustering of samples by catchment, although overlap among groups remained, with ANOSIM indicating weak-to-moderate differentiation in the virome composition among catchments (ANOSIM R = 0.3166, p = 0.0462, Figure 5c). No significance was identified upstream/downstream of identified fish passage barriers (ANOSIM R = −0.0456, p = 0.7039, Figure 5d), or at the species level, although interpretation is limited by the low number of virus-positive īnanga pools (ANOSIM R = 0.3927, p = 0.2253, Figure 5e).

## Discussion

Despite occupying the same river networks, īnanga and Gollum galaxiids had significantly different viromes, suggesting that life-history plays a role in the diversity and abundance of galaxiid viruses. This research represents the first large-scale metatranscriptomic study on how landscape connectivity impacts the viromes of diadromous and river-resident galaxiids in Aotearoa, New Zealand. We investigated the diversity and distribution of fish-associated viruses in two fish species within the same genera but with different life-history strategies, revealing that for fish in the genus *Galaxias*, life history likely plays a role in the composition of their viromes. Furthermore, these results suggest that while some spatial structuring of viral communities may occur among catchments, the overall virome composition remains broadly overlapping across the study region among the isolated river-resident Gollum galaxiids.

The viromes of Gollum galaxiids and īnanga were remarkably different, with only one out of 28 pools of īnanga containing a fish-associated virus, compared to Gollum galaxiids which had 12 of 26 pools with at least one fish-associated virus. This finding complements previous work on īnanga in New Zealand, further highlighted by the fact that the sole virus identified in īnanga was from the family *Coronaviridae* and was the only virus identified previously in any diadromous galaxiid (Grimwood et al., 2023). The viromes of Gollum galaxiids were dominated by the family *Circoviridae* but were further distinguished by only one pool of the 26 having two families of fish-associated viruses, with the remaining 11 having only a single viral family. Many of the top BLAST hits for the Gollum galaxiid pools containing viruses in the family *Circoviridae* were to a circovirus identified from Weddell seal (*Leptonychotes weddellii*) fecal samples, attributed to the consumption of infected fish (Patterson et al., 2021). The circoviruses we identified phylogenetically clustered with other known fish circoviruses, corroborating the fact that this is a fish-associated circovirus. While Gollum galaxiids were found to have the greatest viral abundance and diversity of the two species, viruses were uncommon in both species across the sampling range, yet each species was connected by a virus in the family *Coronaviridae* in different catchments. There was no significant difference identified between species in alpha or beta diversity, likely due to the zero-inflation in the data; however, the striking ecological differences between the two species is undeniable.

Our results indicate that juvenile īnanga were not infected by viruses in the larval marine environment or to any real degree in the freshwater environment, which reinforces previous research on their low viral abundance (Grimwood et al., 2023). Interestingly, none of the juvenile fish of either species had detectable fish-associated viruses. This is noteworthy as both juvenile marine and freshwater fish in other studies have been shown to harbour viruses (Breuil et al., 1991; Grotmol et al., 1997; Munday et al., 1992; Munday & Nakai, 1997; Muroga, 2001; Zhang et al., 2025), with research in other species suggesting that juveniles could harbour a greater viral abundance than adults (Wille et al., 2021). This, however, requires further investigation, as our sample size was relatively limited, and the possibility remains that juvenile īnanga harbour divergent viruses we are unable to identify, or viruses concentrated in tissue we didn’t sample. Notwithstanding, this result coupled with the lack of viruses identified in the post-larval Gollum galaxiids, suggests that the greatest influence on the virome of these two species is the residence time in the environment. As īnanga are predominantly an annual species (McDowall, 1990), their time spent in both marine and freshwater environments is limited, whereas the longer-lived Gollum galaxiid has potentially years to accumulate viruses from the environment.

Given the complete lack of viruses identified in the juveniles of either species, we suggest that viruses may accumulate over time through increased exposure, leading to a greater likelihood of infection with increasing fish age, as opposed to larval or transitional migratory infection. This increased likelihood of viral infection over prolonged or repeated exposure times has been previously identified in a variety of organisms (Benton et al., 2018; Lunn et al., 2019; Molloy et al., 2013) and with parasites in fish (Krkosek et al., 2009; Poulin et al., 2012). It is possible that the longer lifespan of Gollum galaxiids compared to an annual species such as īnanga enables the viruses to persist among populations as contact and transmission rates are extended. As this is the first study on river-resident galaxiids, further research is required to elucidate these findings.

Īnanga and Gollum galaxiids, though they partially overlap in distribution, had significantly different viromes, indicating that galaxiids likely exhibit host specificity in viruses. It is well documented in fish that host specificity drives viral composition, along with other environmental conditions (Costa et al., 2021, 2023; Geoghegan et al., 2021; Grimwood et al., 2023; Perry et al., 2022; Waller et al., 2024). Migratory galaxiids have also been shown to have lower parasite infection levels than non-migratory fish (Poulin et al., 2012), and previous studies on migratory galaxiids, īnanga, and giant kōkopu have shown low and zero fish-associated viral loads, respectively (Grimwood et al., 2023). Alpha diversity at the site and catchment levels were not significantly different, but they were both significantly different in beta diversity, suggesting that viral community composition is indeed host-specific across the region for Gollum galaxiids. Lower viral diversity in diadromous fish than in lacustrine non-migratory conspecifics has been previously identified (Perry et al., 2022). With the additional results from our study and Grimwood et al. (2023), there is an increasing body of evidence that migratory galaxiids may not harbour many fish-specific viruses. Likewise, there were no significant differences in alpha or beta diversity upstream and downstream of the identified fish passage barriers, indicating that within-river connectivity was not an issue. This result further indicates an unidentified medium or vector behind the significant differences in viromes between species and patterns across the region. Further research is needed to identify the medium, or elucidate the dynamics in environmental conditions, host population structure, species community composition, diet or habitat that could be the driving force behind the striking difference between the viromes of these two species.

In this study we sampled only two tissue types, gill and liver, both of which have been shown to be reservoirs and entry sites of viruses in fish (Costa et al., 2024; Kim & Leong, 1999). There is the possibility that galaxiids concentrate viruses in other non-sampled tissues. Future research sampling a broader range of tissues may identify host specificity or tissue tropism, often exhibited by viruses (Watashi & Wakita, 2015). Sampling other tissues in galaxiids could reveal additional viruses not associated with gill and liver tissue. Combining individuals with low viral yields into larger groups may also increase the likelihood of virus detection. However, this would reduce the total number of pools available for analysis, thereby reducing the spatial resolution of the dataset.

Overall, this study expands our understanding of the viromes of galaxiids in Aotearoa, New Zealand, including the first exploration of a river-resident galaxiid. Our findings that īnanga harbour very few fish viruses are surprising but supported by previous research; however, the results on Gollum galaxiids are quite striking. Notably, species that are isolated both genetically and at the population level are the most uniform and abundant in their viromes. While the viromes were dominated by the family *Circoviridae*, four other families of viruses were present across the region. This was contrary to our expectations that migratory īnanga would harbour a more abundant and diverse virome. As this is the first study on the viromes of river-resident galaxiids, it is unknown whether this pattern is consistent throughout the lineage of “roundhead” galaxiids or is unique to Gollum galaxiids. Our study elucidates how fish viromes are shaped by their life history and location. Gaining a better understanding of their viromes can aid in the conservation of these endemic and threatened species, as we learn how the compounding effects of viral load, microbial infections, and environmental conditions/stressors can lead to disease in fish (Darnley et al., 2026). Further work to expand our understanding of galaxiid viromes across species and regions will help us better understand the role of host life history and connectivity in shaping galaxiid viromes as well as the potential transmission pathways through the environment.

## Data accessibility and benefit-sharing

Virus sequences are available on NCBI’s GenBank database under accessions [pending]. The raw reads from fish are available on the Aotearoa Genomics Data Repository [pending].

## Acknowledgements

We would like to thank Polly Bulling, Nuwan DeSilva, Emma Duncan, Roger Hodson and everyone else who assisted us from Environment Southland. To Adam Kitchen, Jeremy Littlejohn, Olivier Raven, Matt Ward and Spencer Kielar for their assistance with fieldwork. To every farmer and sharemilker who welcomed us onto their property to survey the streams that ran through looking for fish, this research was not possible without your generosity.

## Author contributions

Ryan R. Easton: Conceptualisation, performed research, collected samples, laboratory analysis, analysed data, writing.

Stephanie Waller: data analysis, review and editing.

Hayley Bosley: performed research, collected samples, review and editing.

Isa de Vries: laboratory analysis, review.

Matt Jarvis: performed research, collected samples, review.

Jeremy Dubrulle: laboratory analysis, review and editing.

James Hodgkinson-Bean: laboratory analysis, review.

Gerry P. Closs: Conceptualisation, supervision, writing.

Jemma L. Geoghegan: Conceptualisation, supervision, funding, analysed data, writing.

